# Genome sequencing reveals that *Streptococcus pneumoniae* possesses a large and diverse repertoire of antimicrobial toxins

**DOI:** 10.1101/203398

**Authors:** Reza Rezaei Javan, Andries J van Tonder, James P King, Caroline L Harrold, Angela B Brueggemann

**Affiliations:** The Peter Medawar Building for Pathogen Research, Nuffield Department of Medicine, University of Oxford, Oxford, OX1 3SY, United Kingdom.; Division of Infectious Diseases, Department of Medicine, Imperial College London, London, W12 0NN, United Kingdom.

**Keywords:** Bacteriocins, Competition, Antimicrobials, Genomic Epidemiology, Transcriptomics

## Abstract

*Streptococcus pneumoniae* (‘pneumococcus’) is a leading cause of morbidity and mortality worldwide and a frequent coloniser of the nasopharynx. Competition among bacterial members of the nasopharynx is believed to be mediated by bacteriocins: antimicrobial toxins produced by bacteria to inhibit growth of other bacteria. Bacteriocins are also promising candidates for novel antimicrobials. Here, 14 newly-discovered bacteriocin gene clusters were identified among >6,200 pneumococcal genomes. The molecular epidemiology of the bacteriocin clusters was investigated using a large global and historical pneumococcal dataset. The analyses revealed extraordinary bacteriocin diversity among pneumococci and the majority of bacteriocin clusters were also found in other streptococcal species. Genomic hotspots for the integration of bacteriocin genes were discovered. Experimentally, bacteriocin genes were transcriptionally active when the pneumococcus was under stress and when two strains were competing in broth co-culture. These findings fundamentally expand our understanding of bacteriocins relative to intraspecies and interspecies nasopharyngeal competition.

## Introduction

Pneumococci are a leading cause of severe infections such as pneumonia, bacteraemia and meningitis, and are among the most common causes of otitis media, sinusitis and conjunctivitis. All age groups are susceptible to pneumococcal infection, but young children, the elderly, and immunocompromised persons are most at risk (Bogaert et al, 2004). Despite the use of antimicrobials and pneumococcal conjugate vaccines, the pneumococcus remains a major global health problem, causing approximately 14.5 million cases of serious disease and 826,000 deaths annually in children <5 years of age (O’Brien et al., 2009). Antimicrobial-resistant pneumococci have been a serious and increasing concern for several decades (Doern et al., 2001; Klugman, 1990; Tadesse et al., 2017). The pneumococcus is now considered to be a ‘priority pathogen’ – antimicrobial-resistant bacteria that pose the greatest threat to global health – by the World Health Organisation (World Health Organization, 2017).

The pneumococcus is normally an asymptomatic coloniser of the nasopharynx in healthy young children; however, colonisation is also the initial stage of the disease process. The polysaccharide capsule is the main virulence factor of the pneumococcus: nearly 100 distinct capsular antigenic types (serotypes) have been described and certain serotypes are predominantly associated with disease whilst others are largely associated with nasopharyngeal colonisation (Brueggemann et al., 2003). Pneumococci frequently co-colonise with other pneumococci and non-pneumococcal bacterial species, and intraspecies and interspecies competition can influence colonisation dynamics, strain prevalence, serotype distributions and consequently, the potential for disease progression (Shak et al., 2013).

Competition among bacterial members of the nasopharyngeal microbiome is believed to be mediated by bacteriocins: ribosomally-synthesised antimicrobial toxins produced by bacterial species to inhibit the growth of other closely-related bacteria. The producer strain also encodes an immunity protein to protect itself from its own bacteriocin (Czaplewski et al., 2016; Dawid et al., 2006). Bacteriocin production has been associated with more efficient colonisation of a host by the producer strain, owing to the ability of these toxins to remove competitors (Dawid et al., 2006; Shak et al., 2013). Their ability to kill bacteria makes bacteriocins attractive potential candidates for the development of new antimicrobials. Several bacteriocins (e.g. nisin and pediocin PA-1) have already been commercialised and are widely used as food preservatives (Czaplewski et al., 2016).

From a genetic perspective, bacteriocins are found in gene clusters, whereby the genes involved in toxin production, immunity and transport (exporting and processing the toxin peptide) are situated adjacent to each other in the bacterial genome. The *blp* (bacteriocin-like peptides) cluster is the best-characterised bacteriocin in pneumococcus (Bogaardt et al., 2015; Czaplewski et al., 2016; Dawid et al., 2006; Reichmann et al., 2000). Previous work by our group showed that the *blp* cluster is ubiquitous among pneumococci recovered from the early 1900s onward and is highly diverse. We also identified a novel bacteriocin cluster that we named pneumocyclicin (Bogaardt et al., 2015). Five additional bacteriocins have been reported among pneumococci (Begley et al., 2009; Guiral et al., 2005; Hoover et al., 2015; Kadam et al., 2017; Maricic et al., 2016); however, their prevalence, genetic composition and molecular epidemiology in the context of the pneumococcal population are unknown.

The abundance of whole genome sequence data expedites the detection and genetic analysis of bacteriocin clusters. Here, we report 14 newly-discovered pneumococcal bacteriocin clusters and describe the molecular epidemiology of all the currently known bacteriocins within a collection of pneumococci isolated over the past 90 years. We also provide transcriptomic evidence that multiple bacteriocin clusters were induced in response to external stress and also in response to competition for space and nutrients in broth co-culture.

## Results

### Genome mining triples the number of known bacteriocins among pneumococci

Our investigation of a large and diverse dataset of 571 historical and modern pneumococcal genomes resulted in the identification of 14 newly-discovered bacteriocin clusters, increasing the number of known bacteriocins in this species to 21 (Table 1). We identified several clusters similar to lactococcin 972, sactipeptide and lassopeptide bacteriocins (Arnison et al., 2013; Letzel et al., 2014; Martinez et al., 1999), which were hitherto not known to be harboured by the pneumococcus (Table 1; Supplemental Table 1). We subsequently expanded our search for bacteriocins to a much larger dataset of 5,673 published pneumococcal genomes, but no additional bacteriocins were found; thus, the detailed description of the bacteriocins here is restricted to those identified in the dataset of 571 genomes.

**Table 1.** Bacteriocin clusters identified among a dataset of 571 pneumococci recovered since 1916 from patients of all ages residing in 39 different countries.

| Bacteriocin | Genome(s) |  |  | Year(s)<br>of Isolation | Countries<br>(n) | CCs<sup>a</sup><br>(n) | Serotypes<br>(n) |
| --- | --- | --- | --- | --- | --- | --- | --- |
|  | Complete | Partial | Total |  |  |  |  |
| streptococcin A | 404 (70.8%) | 1 (0.2%) | 405 (70.9%) | 1916 - 2009 | 36 | 80 | 76 |
| streptococcin B | 414 (72.5%) | 157 (27.5%) | 571 (100%) | 1916 - 2009 | 39 | 99 | 88 |
| streptococcin C | 571 (100%) | 0 (0.0%) | 571 (100%) | 1916 - 2009 | 39 | 99 | 88 |
| streptococcin D | 6 (1.1%) | 0 (0.0%) | 6 (1.1%) | 1968 - 2005 | 5 | 3 | 4 |
| streptococcin E | 93 (16.3%) | 477 (83.5%) | 570 (99.8%) | 1916 - 2009 | 39 | 98 | 88 |
| streptolancidin A<sup>b</sup> | 1 (0.2%) | 1 (0.2%) | 2 (0.4%) | 1972 - 2006 | 2 | 2 | 2 |
| streptolancidin B<sup>c</sup> | 49 (8.6%) | 44 (7.7%) | 93 (16.3%) | 1939 - 2006 | 16 | 11 | 10 |
| streptolancidin C | 96 (16.8%) | 0 (0.0%) | 96 (16.8%) | 1937 - 2006 | 13 | 15 | 12 |
| streptolancidin D | 49 (8.6%) | 0 (0.0%) | 49 (8.6%) | 1938 - 2006 | 13 | 13 | 11 |
| streptolancidin E<sup>d</sup> | 12 (2.1%) | 156 (27.3%) | 168 (29.4%) | 1937 - 2009 | 26 | 21 | 38 |
| streptolancidin F | 23 (4.0%) | 0 (0.0%) | 23 (4.0%) | 1937 - 2006 | 7 | 4 | 11 |
| streptolancidin G<sup>e</sup> | 190 (33.3%) | 9 (1.6%) | 199 (34.9%) | 1916 - 2009 | 25 | 38 | 46 |
| streptolancidin H | 1 (0.2%) | 0 (0.0%) | 1 (0.2%) | 2006-2008 | 1 | 0 | 1 |
| streptolancidin I | 1 (0.2%) | 0 (0.0%) | 1 (0.2%) | 2009 | 1 | 1 | 1 |
| streptolancidin J | 185 (32.4%) | 195 (34.2%) | 380 (66.5%) | 1916 - 2009 | 33 | 64 | 74 |
| streptolancidin K | 1 (0.2%) | 10 (1.8%) | 11 (1.9%) | 1943 - 2009 | 4 | 2 | 3 |
| streptocyclin<sup>f</sup> | 209 (36.6%) | 0 (0.0%) | 209 (36.6%) | 1937 - 2009 | 20 | 46 | 59 |
| streptolassin | 20 (3.5%) | 0 (0.0%) | 20 (3.5%) | 1939 - 1996 | 8 | 7 | 10 |
| streptosactin | 1 (0.2%) | 0 (0.0%) | 1 (0.2) | 2009 | 1 | 1 | 1 |
| cib<sup>g</sup> | 557 (97.5%) | 0 (0.0%) | 557 (97.5%) | 1916 - 2009 | 39 | 97 | 88 |
| blp<sup>h</sup> | N/A | N/A | 571 (100%) | 1916 - 2009 | 39 | 99 | 88 |

**a.** CC, clonal complex (genetic lineage). Singletons (single genotypes with no closely-related variant) were excluded from the count. Synonym(s) for the previously identified bacteriocins are as follows: **b.** pneumolancidin (Maricic et al., 2016) and salivaricin E (Walker et al., 2016); **c.** lcpAMT (Kadam et al., 2017) and ICESp23FST81 lantibiotic (Croucher et al., 2008); **d.** SP23-BS72 lantibiotic (Begley et al., 2009); **e.** Phr lantibiotic (Hoover et al., 2015); **f.** pneumocyclin (Bogaardt et al., 2015); **g.** *cibAB* (Guiral et al., 2005); **h.** *spi* and *pnc* (Bogaardt et al., 2015).

BLAST searches of the NCBI database of non-redundant nucleotide sequences revealed that the majority of these clusters were not exclusive to pneumococcus, but that identical or similar versions were present in other streptococci (Supplemental Table 2). Therefore, we used the prefix “strepto” and named each cluster according to its type of bacteriocin (see Materials and methods). Among the putative bacteriocins, five were similar to lactococcin 972 (Martinez et al., 1999) and we called these streptococcins (Fig. 1). Despite gene synteny among the streptococcins (Fig. 1A), nucleotide sequence similarity was low: toxin genes, 37-59%; immunity genes, 27-48%; and transporter genes, 49-63% (Fig. 1B). Different streptococcins were found in different, but consistent, locations within the bacterial chromosome (Fig. 1A and 1C). Eleven different streptolancidins, one streptosactin, one streptolassin and one streptocyclicin (what we previously called pneumocyclicin) were also identified among pneumococcal genomes (Fig. 2; Arnison et al., 2013). The bacteriocin clusters were commonly flanked by genes predicted to encode CAAX proteins (possibly involved in self-immunity from bacteriocin toxins; Kjos et al., 2010), Rgg and PlcR transcriptional regulators (implicated in bacterial quorum sensing; Perez-Pascual et al., 2016), transporters, lipoproteins and mobilisation proteins.

**Figure 1.**
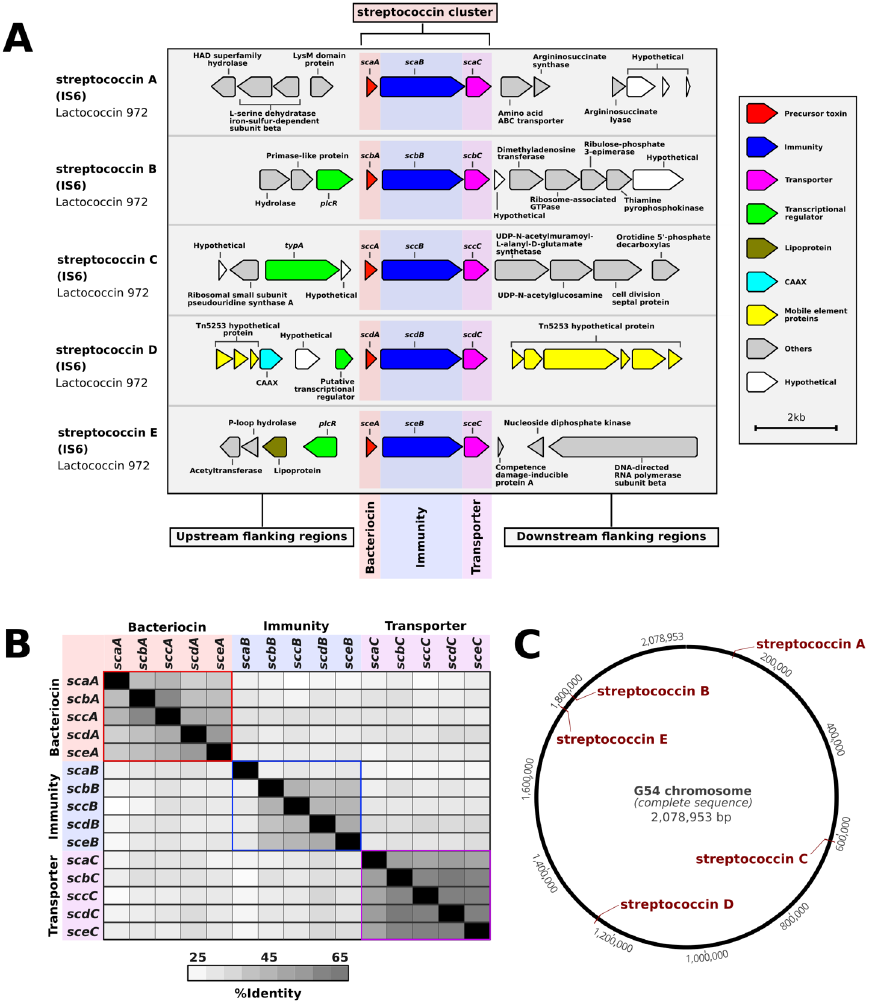
Discovery of streptococcin bacteriocins in the pneumococcus. (A) A schematic representation of each streptococcin cluster and their flanking regions is depicted. The coding regions were derived from the genome of pneumococcal strain IS6, which harbours five different complete streptococcin clusters. (B) A distance matrix of the nucleotide sequence identity shared between genes of different streptococcin clusters found in pneumococcal strain IS6 is shown. (C) Schematic of the finished genome of pneumococcal strain G54 with the locations of the five streptococcin clusters highlighted in red.

**Figure 2.**
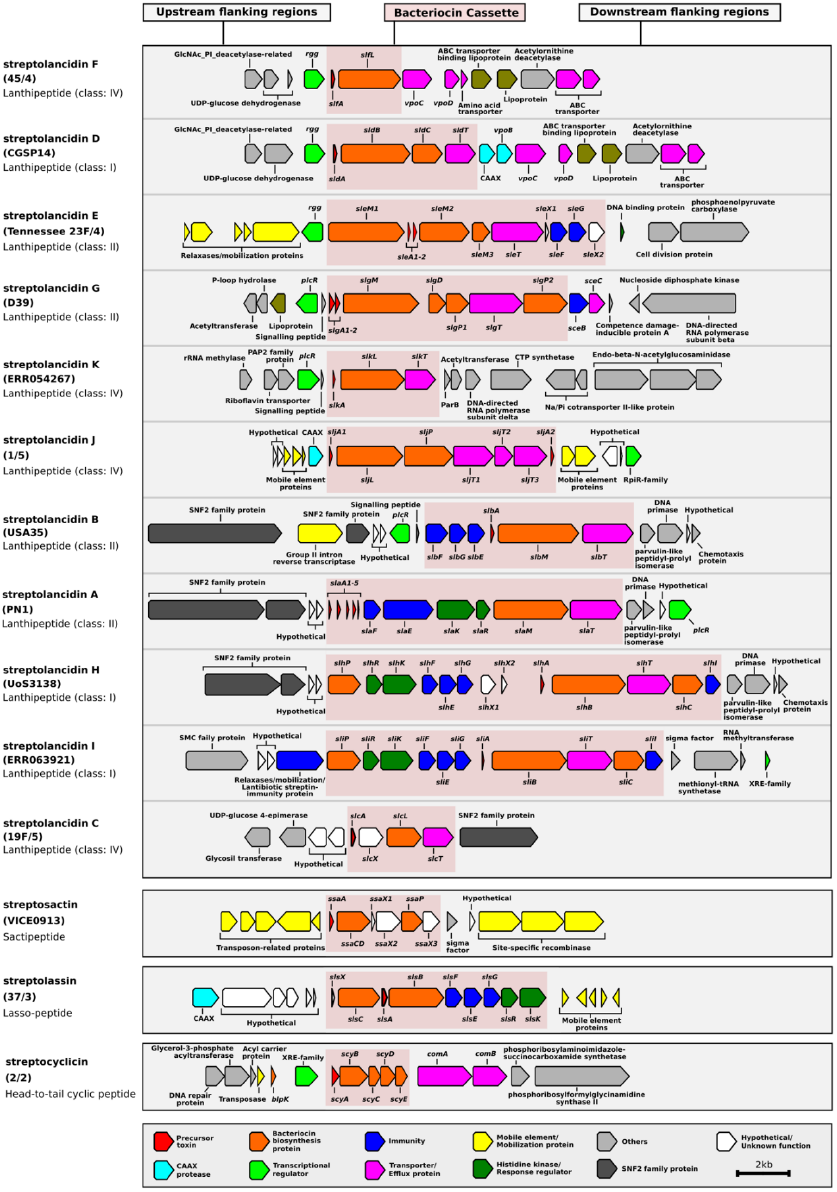
Genetic composition of streptolancidin, streptocyclicin, streptosactin, and streptolassin bacteriocins in the pneumococcal genomes. A schematic representation of 11 different streptolancidins, one streptocyclicin, one streptosactin and one streptolassin identified among pneumococcal genomes is depicted. The names of the isolates containing each bacteriocin cluster are given in brackets and the class of each bacteriocin cluster is given underneath the names of isolates.

Further support for interspecies exchange of bacteriocin gene clusters was provided by the guanine (G) and cytosine (C) content of the bacteriocin clusters. The average GC-content of all pneumococcal genomes in this dataset was 39.6%, whereas the range of values for different bacteriocin groups was as follows: streptococcins, 36.3-42.4%; streptolancidins subset 1, 31.2-33.4%; streptolassin, 31.2%; streptolancidins subset 2, 28.9-29.6%; streptocyclicin, 27.0%; and streptosactin, 25.5% (Supplemental Fig. 1). By comparison, the GC-content of other non-pneumococcal *Streptococcus* species in another study ranged between 33.2-44.6% (Kurioka et al., 2017).

### Extraordinary bacteriocin diversity within a globally-distributed pneumococcal dataset dating from 1916

We further assessed these bacteriocins in the context of the pneumococcal population structure. The study dataset consisted of a diverse collection of 571 pneumococci isolated between 1916 and 2009 from patients and healthy individuals of all ages residing in 39 different countries across six continents. 88 pneumococcal serotypes and 99 different clonal complexes were represented (Table 1; Supplemental Table 3). All bacteriocins detected more than once in the dataset were identified among pneumococci isolated over several decades and from a variety of different countries (Table 1). Some bacteriocins were present in all pneumococci, whilst others were limited to specific clonal complexes (genetic lineages). Bacteriocin clusters missing one or more genes were defined as partial. The percentages of partial and complete clusters varied between different bacteriocins, e.g. streptococcin E was present in all but one pneumococcal genome but was missing one or more genes in the majority (83.5%) of genomes, while streptococcin C was found as a complete cluster in all genomes (Table 1 and Fig. 3A). We constructed a core genome phylogenetic tree of all isolates and labelled each genome according to the presence or absence of each bacteriocin cluster (Supplemental Fig. 2). Overall, we found that the number of bacteriocins within each genome varied from 5 to 11 per genome and that certain combinations of bacteriocins were well represented in the dataset (Fig. 3B, 3C and Supplemental Fig. 2).

**Figure 3.**
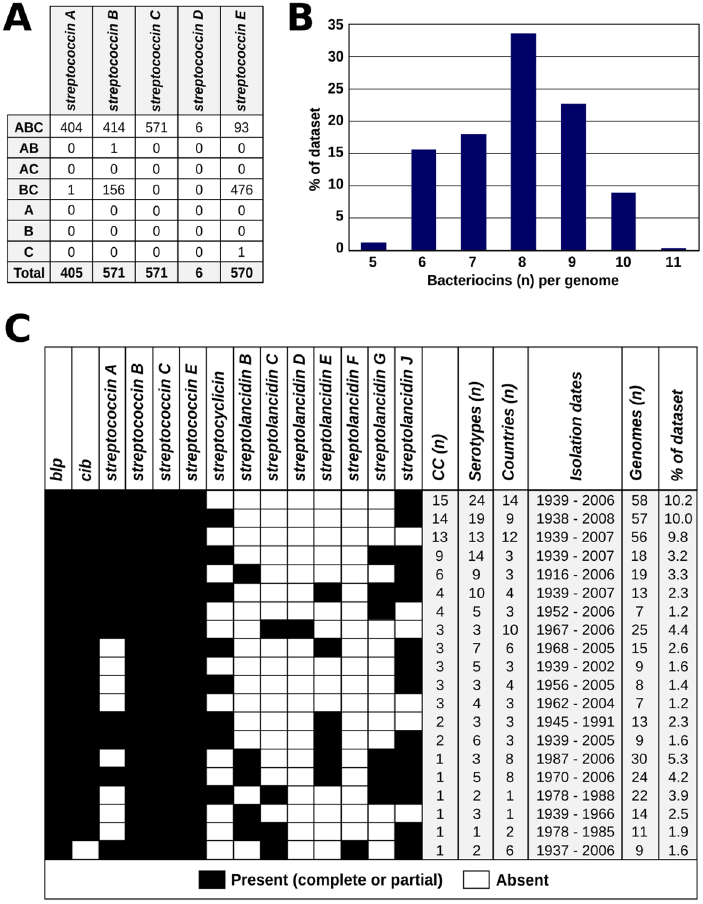
The prevalence, molecular epidemiology and co-occurrence patterns of bacteriocins within the global pneumococcal population. (A) The patterns of gene presence (A, toxin; B, immunity; C, transporter) of different streptococcins within the study dataset are shown. (B) The number of bacteriocins (of any type) per genome among the 571 pneumococcal genomes is summarised. (C) The epidemiological characteristics of the pneumococci that possessed bacteriocin combinations found in >1 percent of the genomes in the dataset are presented.

Due to their relative simplicity (containing only three genes), we chose streptococcins as models for further investigating the pattern of missing genes in partial clusters (Fig. 3A). Scrutinising these clusters revealed that the majority of the partial clusters lack the toxin gene, while still retaining the immunity and/or the transporter genes. This could support the general idea of a ‘cheater’ phenotype, whereby the immunity and transporter genes are conserved to protect the pneumococcus from neighbouring bacteria that express the bacteriocin toxin protein, but the cheater strain does not bear the cost of producing the bacteriocin toxin (Brown et al., 2009; Son et al., 2011).

### Multiple hotspots for the integration of bacteriocin clusters in the pneumococcal genome

By conducting a population genomics-based assessment of the bacteriocin cluster insertion sites, we identified three genomic regions that are putative hotspots for the integration of bacteriocin clusters in the pneumococcal chromosome. These **b**acteriocin **c**luster **h**otspots (BCHs) are specific locations in the genome where different bacteriocin clusters were found in different pneumococci (Fig. 4). This suggested a switching mechanism, whereby different clusters can replace one another via homologous recombination. Up to three different bacteriocin clusters were found to be associated with a single BCH (Fig. 4C). The acquisition of streptolancidin G appears to have rendered the streptococcin E partial by replacing the toxin and part of its immunity gene (Fig. 4A), which is in accordance with the fact that streptolancidin G could not be found in genomes that harboured a complete streptococcin E cluster (Supplemental Fig. 1). Nonetheless, the remnant genes of the streptococcin E partial clusters were conserved in samples collected over many decades (Fig. 3A).

**Figure 4.**
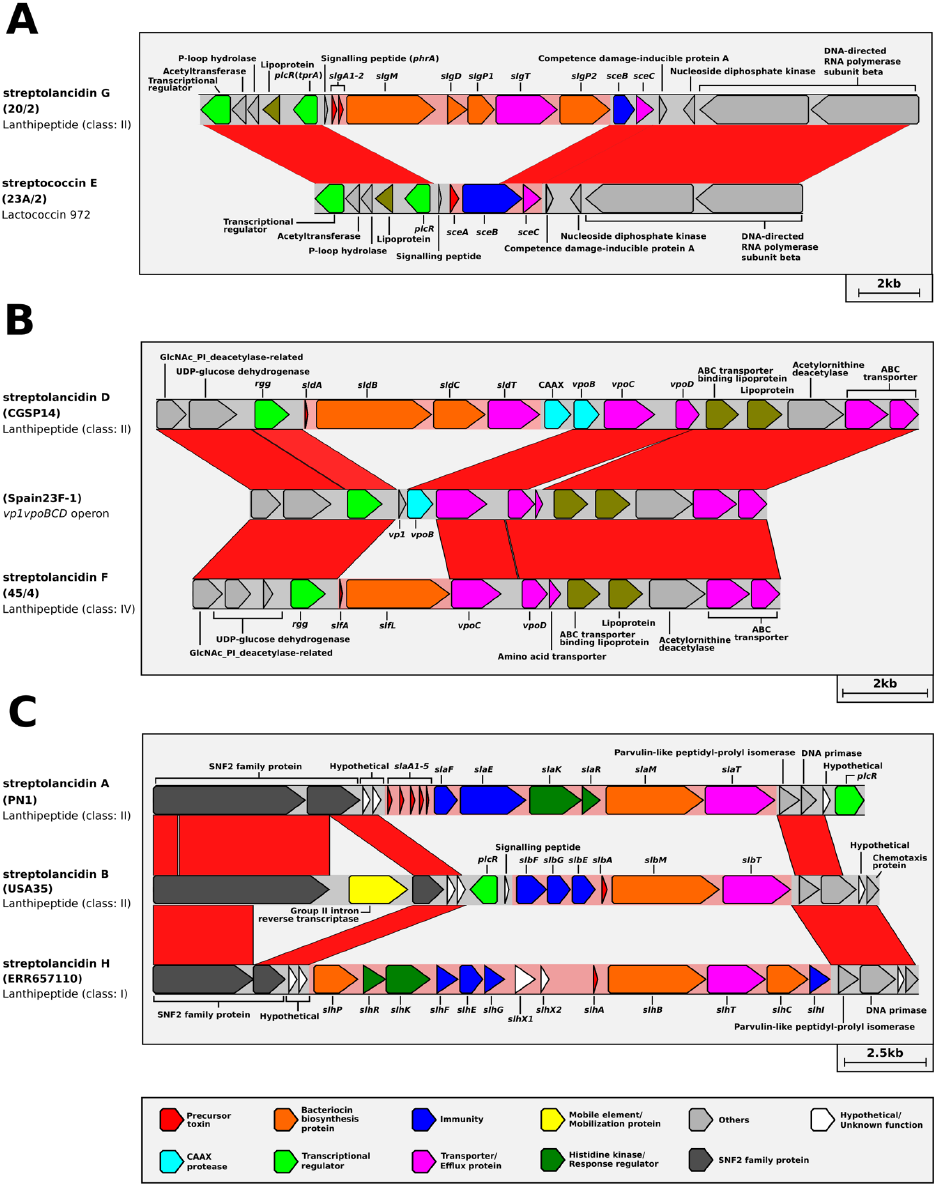
Whole-genome-based population analysis reveals evidence for bacteriocin switching. Bacteriocin cluster hotspots (BCHs) were defined as regions of DNA where different bacteriocin clusters with identical flanking genes were found among pneumococcal genomes. Linear comparisons of (A) BCH-1, (B) BCH-2 and (C) BCH-3 are shown. The isolate names are given in brackets. The class of each bacteriocin cluster is given underneath the isolate name.

### Bacteriocin gene expression in response to heat stress and strain competition

To explore whether the bacteriocin clusters were transcriptionally active, we analysed a whole-genome RNA sequencing dataset from a broth culture of pneumococcal strain 2/2, which was incubated at a higher than normal temperature (40°C vs 37°C) to induce a bacterial stress response (NCBI GEO accession number GSE103778; Kurioka et al., 2017). Multiple genes in the bacteriocin clusters were differentially expressed compared to the control over several time points, indicating that many of these bacteriocin genes were transcribed in response to external stress (Fig. 5).

**Figure 5.**
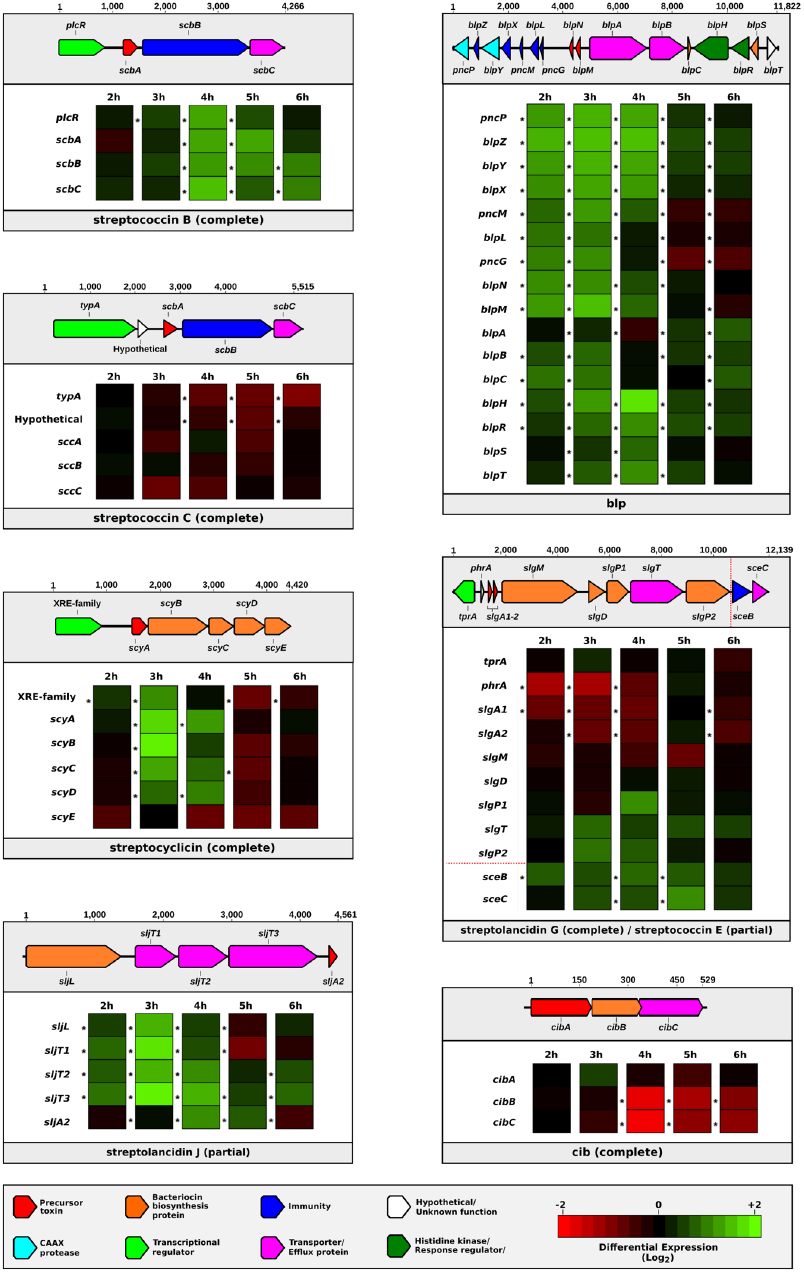
Dynamic changes in the expression of bacteriocin genes in response to bacterial stress. The differential expression levels of bacteriocin gene clusters found in pneumococcal strain 2/2 are displayed in individual boxes. A schematic representation of each bacteriocin cluster is provided above each box. Genes are represented by rows and differential expression levels at different time points are indicated in columns. An asterisk to the left of a cell indicates a statistically significant differential expression level (p <0.05).

Notably, there were two bacteriocin clusters (streptolancidin J and streptococcin E) that were missing genes and yet the remaining genes were clearly being upregulated. Moreover, the timing of gene expression varied across bacteriocin clusters, e.g. genes within a cluster were induced in a specific pattern, whilst at any particular time point during the sampling period several genes in different clusters were upregulated simultaneously. The regulatory governance over which bacteriocin genes are expressed at any given point during bacterial growth remains to be determined.

A second whole-genome RNA sequencing experiment was designed to test whether bacteriocin genes were transcribed when two genetically-different reference strains, PMEN-3 and PMEN-6, were cultured together in the same broth culture using standard incubation conditions. These strains were competing for space and nutrients as the incubation time progressed and cell density increased. Samples were taken for RNA sequencing at multiple time points and sequenced. Sequencing reads from all time points were combined *in silico*. The sequencing reads were mapped to a pseudo-reference genome that was constructed to include the genes unique to PMEN3 and PMEN6 plus the genes shared between both strains (see Materials and methods).

Pneumococcal genes that were significantly upregulated included many genes that would be expected to be expressed during growth (e.g. metabolic genes), in addition to the significant upregulation of 29 bacteriocin genes (Figure 6; Supplemental Table 5). Many of the bacteriocin genes were present in highly similar allelic versions in both PMEN-3 and PMEN-6, thus it was not possible to determine with confidence whether both versions were upregulated or whether one strain overexpressed a similar gene. However, there were three genes within the bacteriocin clusters that were unique: PMEN-3 significantly upregulated the bacteriocin toxin gene of streptococcin A (*scaA*) and a putative immunity gene (*pncM*) from the *blp* bacteriocin cluster, and PMEN-6 significantly upregulated the bacteriocin toxin gene of streptococcin E (*sceA*).

**Figure 6.**
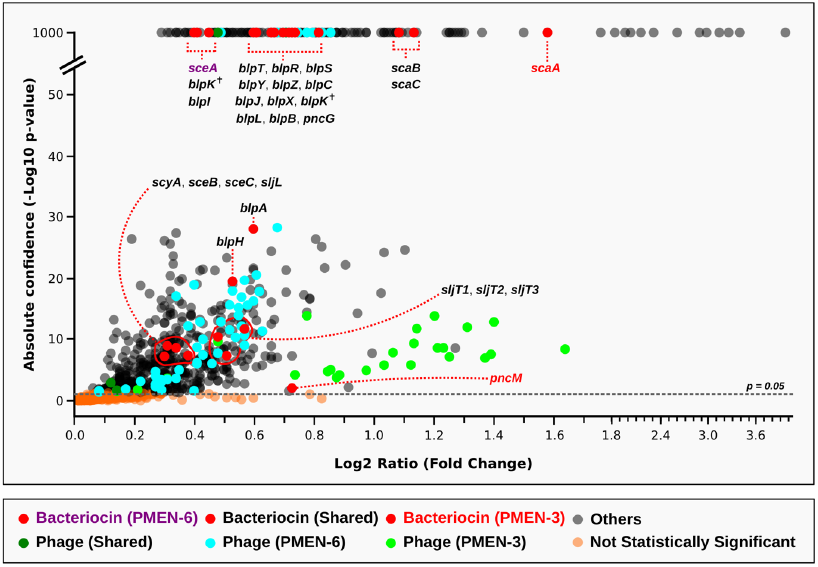
Evidence for the upregulation of bacteriocin genes when two reference strains, PMEN-3 and PMEN-6, were co-cultured in broth media. Only genes that were significantly upregulated compared to the controls (strains cultured individually) are shown in the figure. Genes of interest that were unique to each reference strain and those shared by both strains are marked in different colours. Two copies of *blpK* were found in different locations (one in the *blp* cluster as expected and one elsewhere in the genome) in both genomes and are marked here with a cross. A full list of the genes depicted here and their sequences is found in Supplemental Table 5.

Interestingly, other genes that were significantly upregulated were genes associated with unique prophages present in each of the PMEN strains. We have recently demonstrated that prophage genes are ubiquitous among pneumococcal genomes, but to what extent prophages are influencing pneumococcal biology and perhaps competition between strains is not yet understood (Brueggemann et al., 2017). Overall, the data from this experiment demonstrate proof-of-principle that such methodology can be used to test for the differential expression of key bacterial genes within a competitive environment.

## Discussion

A clear understanding of the role bacteriocins play in pneumococcal biology is central to understanding microbial interactions within the ecological niche (the nasopharynx). The importance of intraspecies competition to pneumococcal ecology is reflected in the changes in prevalence of different pneumococcal serotypes and genotypes in the nasopharynx over time, and understanding competition dynamics is important in the context of understanding vaccine impact (Auranen et al., 2009; Crook et al., 2004; Garcia-Rodriguez et al., 2002). Pneumococcal conjugate vaccines are disruptive to the pneumococcal population structure and alter the composition of microbes competing for space and nutrients in the nasopharynx. The effects of this disruption are not yet fully understood but can lead to increased disease in human populations (Aguiar et al., 2010; Huang et al., 2005; Kaplan et al., 2010).

We revealed here that not only do pneumococci possess a substantially greater and more varied array of bacteriocins than previously recognised, the bacteriocins (often in a particular combination) are associated with specific genetic lineages. This is fundamental, as it provides the framework on which to investigate the mechanisms underpinning specific bacteriocin-pneumococcus combinations, particularly among epidemiologically-successful genetic lineages, and the activity of specific bacteriocin and immunity genes. RNA sequencing clearly demonstrated that bacteriocin genes were transcriptionally active when the pneumococcus was under stress or in competition with another strain during bacterial co-culture. The *in vivo* production of bacteriocin gene products and their functional activities is under further investigation.

Interestingly, we identified several specific locations in the pneumococcal genome that harboured different bacteriocin clusters, which suggested that recombination events had occurred at these locations and resulted in a switching of bacteriocin clusters. The profound impact of recombination on the pneumococcal genome has been described for >25 years and recombination events are well documented in other locations within the pneumococcal genome, most commonly at the capsular polysaccharide locus (conferring a change of serotype) and at penicillin binding protein genes (conferring penicillin resistance if the proteins are altered) (Coffey et al., 1991; Golubchik et al., 2012; Laible et al., 1991; Maynard Smith et al., 1991; Wyres et al., 2012). Similarly, the DpnI, DpnII and DpnIII clusters, each containing a distinct restriction modification system, can replace one another at the *dpn* locus. This is believed to be of protective value in mixed pneumococcal populations against bacteriophages, which are constant invaders of pneumococcal genomes (Brueggemann et al., 2017; Eutsey et al., 2015; Johnston et al., 2013).

We found several examples of presumed bacteriocin cluster switching events that had occurred adjacent to distinct quorum-sensing transcriptional regulators TprA and Rgg. Intriguingly, while the genes that are known to be under the control of TprA and Rgg were replaced, the quorum-sensing transcriptional regulators remained conserved. It is known that TprA controls the expression of its downstream bacteriocin genes: it induces them when pneumococcal cells are at high density in the presence of galactose and represses them when under high glucose growth conditions (Hoover et al., 2015). Galactose is plentiful in the nasopharynx, whereas glucose is scarce (although abundant in the blood), suggesting that the TprA may mediate the expression of its adjacent bacteriocin genes to aid pneumococci to compete for resources during nasopharynx colonisation (Bidossi et al., 2012; Hoover et al., 2015). Likewise, the Rgg quorum-sensing transcriptional regulator has been shown to mediate the expression of its adjacent genes, and this is thought to be directed by sensing amino acid levels in the cellular community (Cuevas et al., 2017). An explanatory hypothesis might be that bacteriocin cluster switching provides a mechanism by which the existing intricate quorum-sensing signalling network required for coordinating population-level behaviours is accessible by the newly-acquired bacteriocin cluster. This remains to be experimentally verified.

Our current work is significant for the broader community in that among the 21 different pneumococcal bacteriocin clusters now identified, many homologues are found in other unrelated streptococcal species. We also provide here a unified nomenclature for the pneumococcal bacteriocin clusters and their genes. Finally, the fact that any single pneumococcal genome possesses multiple bacteriocin clusters should be carefully considered when designing laboratory experiments aimed at assessing the activity of an individual bacteriocin.

Overall, these population genomic and transcriptomic analyses reveal an extraordinary complexity of bacteriocins among pneumococci and underscore the need to determine precisely how these bacteriocins drive changes within the pneumococcal population and the wider microbial community. Such findings are interesting not only for their population biology and ecology insights, but also because bacteriocins potentially have a huge impact on public health. By directly influencing changes in microbial populations, bacteriocins might indirectly be affecting the effectiveness of vaccines in the longer term: after vaccine use in human populations, the target pneumococcal population is significantly changed and those pneumococci must now compete within the altered microbiome. If bacteriocins are essential to the pneumococcal competitive strategy, then the composition of bacteriocins possessed by the post-vaccination pneumococcal community is important to understand. Moreover, there is obvious potential for the development of bacteriocins as novel antimicrobials: at a time when the challenges of antimicrobial-resistant microbes have never been more acute, these data provide many new areas of investigation.

## Materials and methods

### Genome mining for bacteriocin clusters

In total, 6,244 assembled pneumococcal genomes from studies previously published by us and others (Brueggemann et al., 2017; Chewapreecha et al., 2014; Croucher et al., 2011; Croucher et al., 2013a; Croucher et al., 2013b; Gladstone et al., 2015; van Tonder et al., 2014; van Tonder et al. 2015) (Supplemental Table 4) were screened for the presence of bacteriocin clusters using a variety of bioinformatic tools and databases, including antiSMASH (Weber et al., 2015), BACTIBASE (Hammami et al., 2010), BAGEL (de Jong et al., 2006) and InterProScan (Jones et al., 2014) (data collected in March 2016). An in-house pipeline was developed to automate part of this process. Predicted gene clusters from each of the database outputs were examined manually and further scrutinised using extensive BLAST searches.

### Analyses of the putative bacteriocin genes

Putative bacteriocin genes were annotated using homology to other known bacteriocin genes (not shown) in all available databases mentioned above, as well as structure-based searches. Protein domains were examined using the Conserved Domain search feature at NCBI (Marchler-Bauer et al., 2010). Genes of interest were screened against the STRING database (Szklarczyk et al., 2015) to search for any previously reported relationship to other genes. Multiple sequence alignments of the genes in streptococcin clusters were performed in Geneious version 9.1 (Biomatters Ltd) using the ClustalW algorithm (Larkin et al., 2007) with default parameters (Gap open cost=15, Gap extend cost=6.66). The multiple sequence alignment output was used within the Geneious environment to calculate percentage identity matrices. The figures of the coding regions of the bacteriocin clusters and their flanking genes were generated in Geneious and edited using Inkscape (http://inkscape.org).

### Classification and nomenclature of bacteriocin clusters

Putative bacteriocin clusters were classified based on their predicted biosynthetic machinery and structural features (Arnison et al., 2013) and their corresponding genes were designated per the standards of nomenclature for bacteriocins (Supplemental Table 1). BLAST searches in the NCBI database revealed that the majority of these clusters are also present in other closely related streptococci (Supplemental Table 2); therefore, the clusters were named using the prefix “strepto-” followed by an abbreviation of their bacteriocin class: “streptococcins” for those that were lactococcin 972-like, “streptolancidins” for lanthipeptides, “streptocyclicins” for the head-to-tail cyclized peptides, “streptosactins” for the sactipeptides and “streptolassins” for the lassopeptide group of bacteriocins. When more than one bacteriocin cluster from the same class was present, they were lettered alphabetically by the order of their discovery in our analyses.

### Molecular epidemiology of the bacteriocin clusters

We compiled a global and historical dataset (n=571) by selecting a diverse set of pneumococcal genomes isolated between 1916 and 2009 from people of all ages residing in 39 different countries. Samples from both carriage and disease, 88 pneumococcal serotypes and 99 different clonal complexes were represented in this dataset (Supplemental Table 3). Genomes and their associated metadata were stored in a BIGSdb database (Jolley et al., 2010). The BIGSdb database platform was used to generate a presence/absence matrix of all the known bacteriocin genes in all 571 genomes. (Note that there were two small frameshifted gene remnants of both streptolancidins C and E present in some genomes, but these were not analysed further in this study.) Using this matrix (79,940 genes) as an input, an in-house python script was developed (available upon request) to calculate the prevalence, molecular epidemiology and co-occurrence patterns of all the bacteriocin clusters in the study dataset.

### Construction of the core genome phylogenetic tree

All genomes in the study dataset were annotated using the Prokka prokaryotic annotation pipeline (Seemann 2014). The annotation files were input into Roary (Page et al., 2015) and clustered using a sequence identity threshold of 90%. Genes present in every genome were selected using a core genome threshold of 100% and were aligned using Roary. FastTreeMP (Price et al., 2010) was used to construct the phylogenetic tree using generalised time-reversible model (parameters: FastTreeMP -nt -gtr). ClonalFrameML (Didelot et al., 2015) was then applied to reconstruct the phylogenetic tree adjusted for recombination. The tree was annotated using iTOL (Letunic et al., 2016) and Inkscape.

### Investigation of bacteriocin cluster insertion sites

Bacteriocin cluster sequences were used as queries to BLAST against genomes in the study dataset using the custom BLAST implemented in Geneious. The matching region plus additional flanking regions were visualised using the query-centric alignment feature within the Geneious environment. Regions of DNA with different bacteriocin clusters but identical flanking genes among different isolates were identified and further investigated using the Artemis Comparison Tool (ACT) (Carver et al., 2005). Linear comparison figures were generated using Geneious, ACT and Inkscape.

### RNA sequencing analyses

In the first experiment, total bacterial RNA sequencing was performed on RNA extracted from pneumococcal strain 2/2 grown at a higher incubation temperature than normal (40 vs 37°C) to induce a bacterial stress response (Kurioka et al., 2017). Pneumococci were grown in brain-heart infusion broth for 6 h and RNA extractions were performed on samples from five time points (2, 3, 4, 5 and 6 h of incubation) using the Promega Maxwell^®^ 16 Instrument and LEV simplyRNA Cells purification kit, following the manufacturer’s protocol. Extracted RNA samples were sent to the Oxford Genomics Centre for sequencing on the Illumina platform (NCBI GEO accession number GSE103778). The sequenced forward and reverse reads were paired and mapped onto the annotated pneumococcal strain 2/2 genome using Bowtie2 (Langmead et al., 2012) with the highest sensitivity option. Differential gene expression was assessed in Geneious using the DESeq (Anders et al., 2010) method. Genes with an adjusted P value <0.05 were deemed to be differentially expressed.

In a second experiment, pneumococcal reference strains PMEN-3 (Spain^9V^-3) and PMEN-6 (Hungary^19A^-6) were grown together in brain-heart infusion broth for 6 hours. The controls were prepared by growing each strain individually in brain-heart infusion broth for 6 h. Total bacterial RNA sequencing was performed on RNA extracted from broth cultures at 2, 3, 4, 5, and 6 h after incubation using the procedures described above (NCBI GEO accession number GSE110750).

A pseudo-reference genome was constructed using Bowtie2, Velvet (Zerbino et al., 2008), and MeDuSa (Bosi et al., 2015) to sort genes into three categories: those unique to PMEN-3, those unique to PMEN-6, and those shared between the two (Supplemental Fig. 3A). The RNA sequencing reads for each individual control strain at all time points were pooled *in silico*. Data from all time points were combined to minimise possible variability caused by different growth rates of strains. Data from all time points sampled in the broth culture of PMEN-3 + PMEN-6 were also combined *in silico* and mapped to the pseudo-reference genome using Bowtie2 with the highest sensitivity option. Differential expression analyses were performed using the DESeq method by comparing sequence reads generated when strains were co-cultured to those from when strains were grown individually (Supplemental Fig. 3B).

Theoretically, the control contained double the amount of reads in comparison to the *in vivo* competition experiment due to being compiled *in silico* from two sets of samples. While this meant that the downregulation of genes could not reliably be assessed, one could have confidence that upregulated genes were differentially expressed (since expression levels must exceed that of the combined controls); however, this approach can provide only relative and not absolute values, and true fold-change ratios for the upregulated genes are most likely underestimated using this method.

## Data access

Representative examples of the newly-discovered bacteriocin clusters have been deposited at GenBank under the accession numbers of MF990778-MF990796. Accession numbers for all genomes used in this study are listed in Supplemental Table 4. Raw transcriptomic sequence data used in this study is available under GEO accession numbers GSE103778 and GSE110750.

## Acknowledgements

The authors wish to acknowledge computational assistance and helpful comments on the manuscript from Dr Melissa Jansen van Rensburg.

---

**Supplemental Figure 1.**
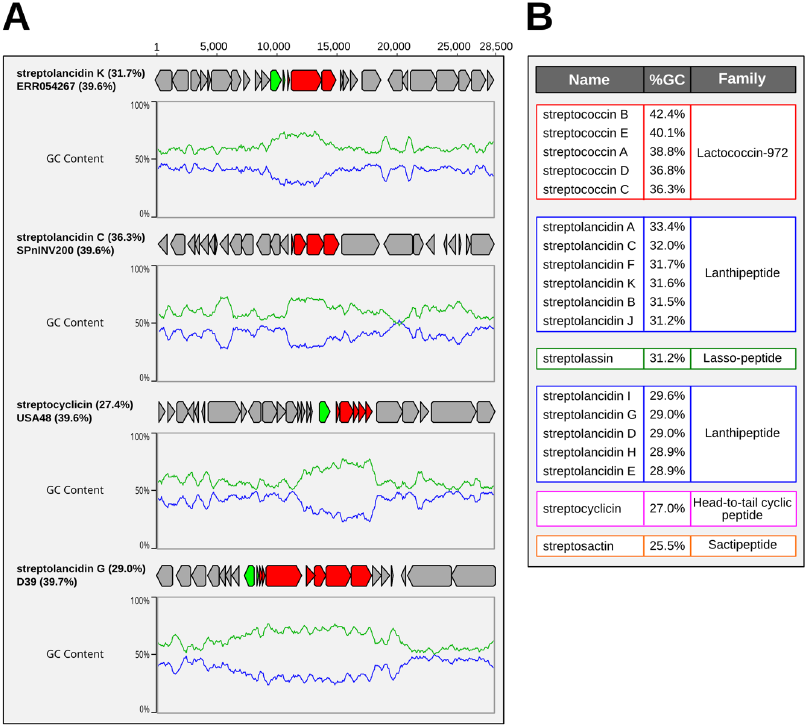
Guanine (G) and cytosine (C) content of pneumococcal bacteriocin clusters. A. Four examples of GC plots depicting the percentage GC-content of bacteriocin cluster genes (red), transcriptional regulator genes (green) and other adjacent pneumococcal genes (grey). The names of the bacteriocin and the genome in which it was found are given, with the percentage GC-content of each in brackets. Each graph depicts GC-content and adenine (A) thymine (T) content by the green and blue lines, respectively. B. Average GC-content for each bacteriocin cluster type, organised by bacteriocin type or class. The lanthipeptides formed two subsets based on GC-content of <30% or >31%.

**Supplementary Figure 2.**
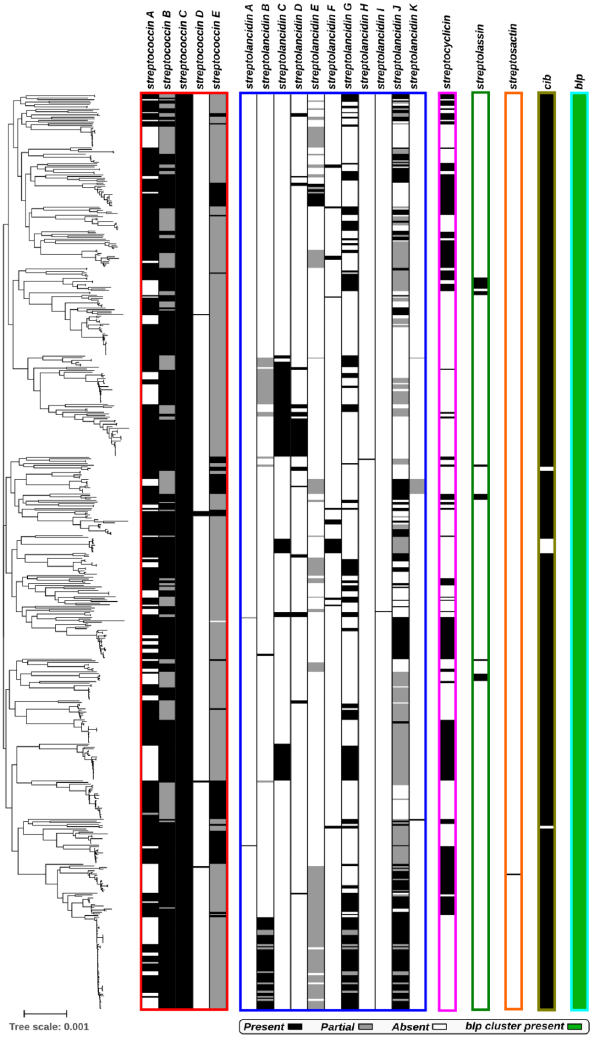
Diversity of bacteriocins within a global pneumococcal dataset. A phylogenetic tree of all genomes in the study dataset is depicted and labelled according to the presence of different bacteriocins. Clusters with missing genes were defined as partial. The exceptions were the *blp* clusters: their highly diverse and complicated genetic compositions among pneumococci genomes meant that a similar classification between partial and complete clusters could not be applied (ref. 12). Instead, their presence (irrespective of being partial or complete) is depicted by the green colour.

**Supplemental Figure 3.**
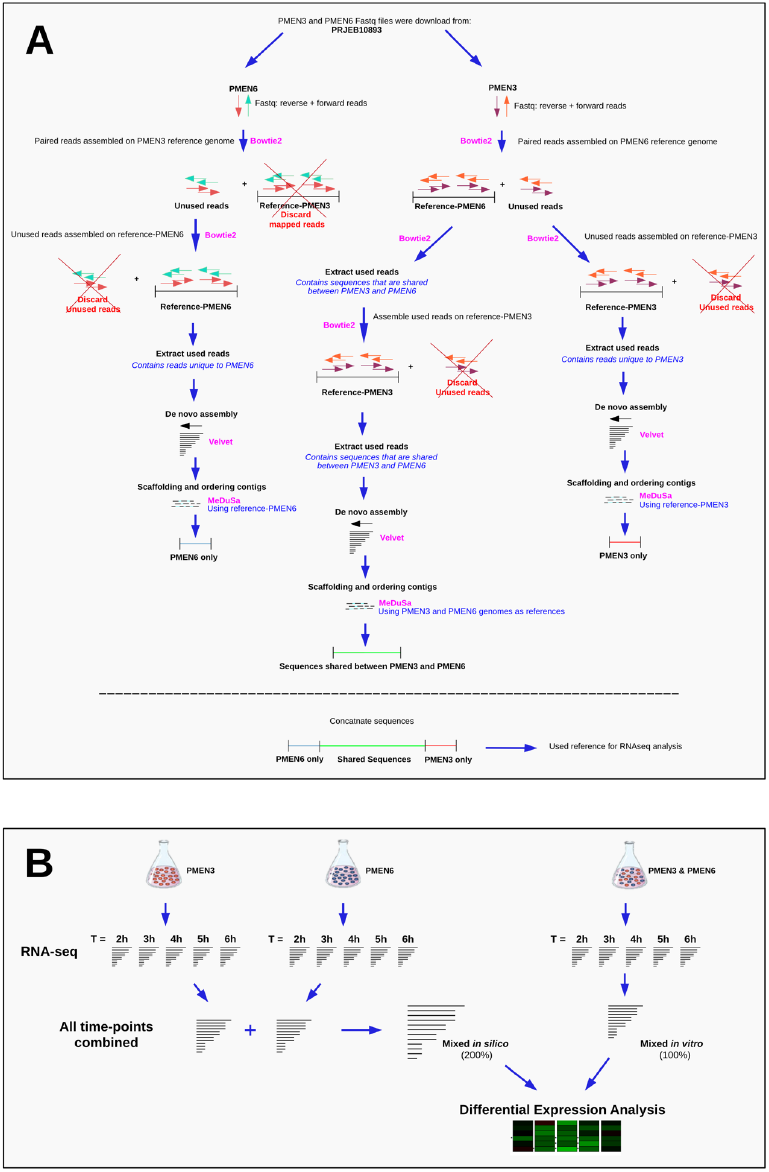
Methodology for analysing the RNA sequencing data from the co-colonisation experiment. A. Steps involved in creating the pseudo-reference genome sequence. The name of the tool used in each step is shown in pink. B. Schematic describing the combination of RNA sequence reads to assess differential expression levels.

## References

1. Aguiar S, Brito M, Gonçalo-Marques J, Melo-Cristino J, Ramirez M. (2010) Serotypes 1, 7F and 19A became the leading causes of pediatric invasive pneumococcal infections in Portugal after 7 years of heptavalent conjugate vaccine use. Vaccine 28: 5167–5173. DOI:10.1016/j.vaccine.2010.06.008

2. Anders S, Huber W. (2010) Differential expression analysis for sequence count data. Genome Biology 11:R106 https://doi.org/10.1186/gb-2010-11-10-r106

3. Arnison P, Bibb M, Bierbaum G, Bowers A, Bugni T, Bulaj G, Camarero JA, Campopiano DJ, Challis GL, Clardy J, Cotter PD, Craik DJ, Dawson M, Dittmann E, Donadio S, Dorrestein PC, Entian KD, Fischbach MA, Garavelli JS, Göransson U, Gruber CW, Haft DH, Hemscheidt TK, Hertweck C, Hill C, Horswill AR, Jaspars M, Kelly WL, Klinman JP, Kuipers OP, Link AJ, Liu W, Marahiel MA, Mitchell DA, Moll GN, Moore BS, Müller R, Nair SK, Nes IF, Norris GE, Olivera BM, Onaka H, Patchett ML, Piel J, Reaney MJ, Rebuffat S, Ross RP, Sahl HG, Schmidt EW, Selsted ME, Severinov K, Shen B, Sivonen K, Smith L, Stein T, Süssmuth RD, Tagg JR, Tang GL, Truman AW, Vederas JC, Walsh CT, Walton JD, Wenzel SC, Willey JM, van der Donk WA. (2013) Ribosomally synthesized and post-translationally modified peptide natural products: overview and recommendations for a universal nomenclature. Nat Prod Rep 30: 108–160. DOI:10.1039/c2np20085f

4. Auranen K, Mehtälä J, Tanskanen A, S. Kaltoft M. (2009) Between-strain competition in acquisition and clearance of pneumococcal carriage–epidemiologic evidence from a longitudinal study of day-care children. Am J Epidemiol 171: 169–176. https://doi.org/10.1093/aje/kwp351

5. Begley M, Cotter P, Hill C, Ross R. (2009) Identification of a novel two-peptide lantibiotic, Lichenicidin, following rational genome mining for LanM proteins. Appl Environ Microbiol 75: 5451–5460. DOI:10.1128/AEM.00730-09

6. Bidossi A, Mulas L, Decorosi F, Colomba L, Ricci S, Pozzi G, Deutscher J, Viti C, Oggioni MR. (2012) A functional genomics approach to establish the complement of carbohydrate transporters in *Streptococcus pneumoniae*. PLOS ONE 7: e33320. https://doi.org/10.1371/journal.pone.0033320

7. Bogaardt C, van Tonder A, Brueggemann AB. (2015) Genomic analyses of pneumococci reveal a wide diversity of bacteriocins – including pneumocyclicin, a novel circular bacteriocin. BMC Genomics 16: 554..DOI:10.1186/s12864-015-1729-4.

8. Bogaert D, de Groot R, Hermans P. (2004) *Streptococcus pneumoniae* colonisation: the key to pneumococcal disease. Lancet Infect Dis 4: 144–154. DOI:10.1016/S1473-3099(04)00938-7

9. Bosi E, Donati B, Galardini M, Brunetti S, Sagot M, Lió P, Crescenzi P, Fani R, Fondi M. (2015) MeDuSa: a multi-draft based scaffolder. Bioinformatics 31: 2443–2451. DOI:10.1093/bioinformatics/btv171

10. Brown S, West S, Diggle S, Griffin A. (2009) Social evolution in micro-organisms and a Trojan horse approach to medical intervention strategies. Phil Trans Royal Soc B 364: 3157–3168. DOI:10.1098/rstb.2009.0055

11. Brueggemann AB, Griffiths D, Meats E, Peto T, Crook D, Spratt B. (2003) Clonal relationships between invasive and carriage S*treptococcus pneumoniae* and serotype- and clone-specific differences in invasive disease potential. J Infect Dis 187: 1424–1432. DOI:10.1086/374624

12. Brueggemann AB, Harrold C, Rezaei Javan R, van Tonder A, McDonnell A, Edwards B. (2017) Pneumococcal prophages are diverse, but not without structure or history. Sci Rep 7: 42976. DOI:10.1038/srep42976

13. Carver T, Rutherford K, Berriman M, Rajandream M, Barrell B, Parkhill J. (2005) ACT: the Artemis comparison tool. Bioinformatics 21: 3422–3423. DOI:10.1093/bioinformatics/bti553

14. Chewapreecha C, Harris S, Croucher N, Turner C, Marttinen P, Cheng L, Pessia A, Aanensen DM, Mather AE, Page AJ, Salter SJ, Harris D, Nosten F, Goldblatt D, Corander J, Parkhill J, Turner P, Bentley SD. (2014) Dense genomic sampling identifies highways of pneumococcal recombination. Nat Genet 46: 305–309. DOI:10.1038/ng.2895

15. Coffey T, Dowson C, Daniels M, Zhou J, Martin C, Spratt B, Musser JM. (1991) Horizontal transfer of multiple penicillin-binding protein genes, and capsular biosynthetic genes, in natural populations of *Streptococcus pneumoniae*. Mol Microbiol 5: 2255–2260. PMID: 1766389

16. Crook DW, Brueggemann AB, Sleeman K, Peto TEA. (2004) Pneumococcal carriage. In The Pneumococcus (eds. Tuomanen EI, Mitchell TJ, Morrison DA, Spratt B.) 136–147 (ASM Press, Washington (D.C.), USA).

17. Croucher N, Finkelstein J, Pelton S, Mitchell P, Lee G, Parkhill J, Bentley SD, Hanage WP, Lipsitch M. (2013) Population genomics of post-vaccine changes in pneumococcal epidemiology. Nat Genet 45: 656–663. DOI:10.1038/ng.2625

18. Croucher N, Harris S, Fraser C, Quail M, Burton J, van der Linden M, McGee L, von Gottberg A, Song JH, Ko KS, Pichon B, Baker S, Parry CM, Lambertsen LM, Shahinas D, Pillai DR, Mitchell TJ, Dougan G, Tomasz A, Klugman KP, Parkhill J, Hanage WP, Bentley SD. (2011) Rapid pneumococcal evolution in response to clinical interventions. Science 331: 430–434. DOI:10.1126/science.1198545

19. Croucher N, Mitchell A, Gould K, Inverarity D, Barquist L, Feltwell T, Fookes MC, Harris SR, Dordel J, Salter SJ, Browall S, Zemlickova H, Parkhill J, Normark S, Henriques-Normark B, Hinds J, Mitchell TJ, Bentley SD. (2013) Dominant role of nucleotide substitution in the diversification of serotype 3 pneumococci over decades and during a single infection. PLOS Genet 9: e1003868. DOI:10.1371/journal.pgen.1003868

20. Croucher N, Walker D, Romero P, Lennard N, Paterson G, et al. (2008) Role of conjugative elements in the evolution of the multidrug-resistant pandemic clone *Streptococcus pneumoniae* Spain23F-ST81. J Bacteriol 191: 1480–1489. DOI:10.1128/JB.01343-08

21. Cuevas R, Eutsey R, Kadam A, West-Roberts J, Woolford C, Mitchell AP, Mason KM, Hiller NL. (2017) A novel streptococcal cell-cell communication peptide promotes pneumococcal virulence and biofilm formation. Mol Microbiol 105: 554–571. DOI:10.1111/mmi.13721

22. Czaplewski L, Bax R, Clokie M, Dawson M, Fairhead H, Fischetti VA, Foster S, Gilmore BF, Hancock RE, Harper D, Henderson IR, Hilpert K, Jones BV, Kadioglu A, Knowles D, Ólafsdóttir S, Payne D, Projan S, Shaunak S, Silverman J, Thomas CM, Trust TJ, Warn P, Rex JH. (2016) Alternatives to antibiotics—a pipeline portfolio review. Lancet Infect Dis 16: 239–251. DOI:10.1016/S1473-3099(15)00466-1

23. Dawid S, Roche A, Weiser J. (2006) The *blp* bacteriocins of *Streptococcus pneumoniae* mediate intraspecies competition both *in vitro* and *in vivo*. Infect Immun 75: 443–451. DOI:10.1128/IAI.01775-05

24. Didelot X, Wilson D. (2015) ClonalFrameML: efficient inference of recombination in whole bacterial genomes. PLoS Comput Biol. 11: e1004041. DOI:10.1371/journal.pcbi.1004041

25. Doern G, Heilmann K, Huynh H, Rhomberg P, Coffman S, Brueggemann AB. (2001) Antimicrobial resistance among clinical isolates of *Streptococcus pneumoniae* in the United States during 1999-2000, including a comparison of resistance rates since 1994-1995. Antimicrob Agents Chemother 45: 1721–1729. DOI:10.1128/AAC.45.6.1721-1729.2001

26. Eutsey R, Powell E, Dordel J, Salter S, Clark T, Clark TA, Korlach J, Ehrlich GD, Hiller NL. (2015) Genetic stabilization of the drug-resistant PMEN1 pneumococcus lineage by its distinctive DpnIII restriction-modification system. mBio 6: e00173–15. DOI:10.1128/mBio.00173-15

27. Garcia-Rodriguez J, Fresnadillo Martínez M. (2002) Dynamics of nasopharyngeal colonization by potential respiratory pathogens. J Antimicrob Chemother 50: 59–74. https://doi.org/10.1093/jac/dkf506

28. Gladstone R, Jefferies J, Tocheva A, Beard K, Garley D, Chong WW, Bentley SD, Faust SN, Clarke SC. (2015) Five winters of pneumococcal serotype replacement in UK carriage following PCV introduction. Vaccine 33: 2015–2021. DOI:10.1016/j.vaccine.2015.03.012

29. Golubchik T, Brueggemann AB, Street T, Gertz R, Spencer C, Ho T, Giannoulatou E, Link-Gelles R, Harding RM, Beall B, Peto TE, Moore MR, Donnelly P, Crook DW, Bowden R. (2012) Pneumococcal genome sequencing tracks a vaccine escape variant formed through a multi-fragment recombination event. Nat Genet 44: 352–355. DOI:10.1038/ng.1072

30. Guiral S, Mitchell T, Martin B, Claverys J. (2005) Competence-programmed predation of noncompetent cells in the human pathogen *Streptococcus pneumoniae*: genetic requirements. Proc Natl Acad Sci USA 102: 8710–8715. https://doi.org/10.1073/pnas.0500879102

31. Hammami R, Zouhir A, Le Lay C, Ben Hamida J, Fliss I. (2010) BACTIBASE second release: a database and tool platform for bacteriocin characterization. BMC Microbiol 10: 22. DOI:10.1186/1471-2180-10-22

32. Hoover S, Perez A, Tsui H, Sinha D, Smiley D, DiMarchi RD, Winkler ME, Lazazzera BA. (2015) A new quorum-sensing system (TprA/PhrA) for *Streptococcus pneumoniae* D39 that regulates a lantibiotic biosynthesis gene cluster. Mol Microbiol 97: 229–243. DOI:10.1111/mmi.13029

33. Huang S, Platt R, Rifas-Shiman S, Pelton S, Goldmann D, Finkelstein J. (2005) Post-PCV7 changes in colonizing pneumococcal serotypes in 16 Massachusetts communities, 2001 and 2004. Pediatrics 116: e408–e413. www.pediatrics.org/cgi/doi/.10.1542/peds.2004-2338

34. Johnston C, Polard P, Claverys J. (2013) The DpnI/DpnII pneumococcal system, defense against foreign attack without compromising genetic exchange. Mob Genet Elements 3: e25582. DOI:10.4161/mge.25582

35. Jolley K, Maiden M. (2010) BIGSdb: Scalable analysis of bacterial genome variation at the population level. BMC Bioinform 11: 595. DOI:10.1186/1471-2105-11-595

36. Jones P, Binns D, Chang H, Fraser M, Li W, McAnulla C, McWilliam H, Maslen J, Mitchell A, Nuka G, Pesseat S, Quinn AF, Sangrador-Vegas A, Scheremetjew M, Yong SY, Lopez R, Hunter S. (2014). InterProScan 5: genome-scale protein function classification. Bioinformatics 30: 1236–1240. DOI:10.1093/bioinformatics/btu031

37. Kadam A, Eutsey R, Rosch J, Miao X, Longwell M, Xu W, Woolford CA, Hillman T, Motib AS, Yesilkaya H, Mitchell AP, Hiller NL. (2017) Promiscuous signaling by a regulatory system unique to the pandemic PMEN1 pneumococcal lineage. PLOS Pathog 13: e1006339. https://doi.org/10.1371/journal.ppat.1006339

38. Kaplan S, Barson W, Lin P, Stovall S, Bradley J, Tan TQ, Hoffman JA, Givner LB, Mason EO Jr. (2010) Serotype 19A is the most common serotype causing invasive pneumococcal infections in children. Pediatrics 125: 429–436. DOI:10.1542/peds.2008-1702

39. Kjos M, Snipen L, Salehian Z, Nes I, Diep D. (2010) The Abi proteins and their involvement in bacteriocin self-immunity. J Bacteriol 192: 2068–2076. DOI:10.1128/JB.01553-09

40. Klugman K. (1990) Pneumococcal resistance to antibiotics. Clin Microbiol Rev 3: 171–196. PMCID: PMC358150

41. Kurioka A, van Wilgenburg B, Javan Rezaei R, Hoyle R, van Tonder AJ, Harrold CL, Leng T, Howson LJ, Shepherd D, Cerundolo V, Brueggemann AB, Klenerman P. (2017) Diverse *Streptococcus pneumoniae* strains drive a MAIT cell response through MR1-dependent and cytokine-driven pathways. J Infect Dis 2017: doi:https://doi.org/10.1093/infdis/jix647

42. Laible G, Spratt BG, Hakenbeck R. (1991) Interspecies recombinational events during the evolution of altered PBP 2x genes in penicillin-resistant clinical isolates of *Streptococcus pneumoniae*. Mol Microbiol 5: 1993–2002. DOI:10.1111/j.1365-2958.1991.tb00821.x

43. Langmead B, Salzberg S. (2012) Fast gapped-read alignment with Bowtie 2. Nat Methods 9: 357–359. DOI:10.1038/nmeth.1923

44. Larkin MA, Blackshields G, Brown NP, Chenna R, McGettigan PA, McWilliam H, Valentin F, Wallace IM, Wilm A, Lopez R, Thompson JD, Gibson TJ, Higgins DG. (2007) Clustal W and Clustal X version 2.0. Bioinformatics 23:2947–8. DOI:10.1093/bioinformatics/btm404

45. Letunic I, Bork P. (2016) Interactive tree of life (iTOL) v3: an online tool for the display and annotation of phylogenetic and other trees. Nucleic Acids Res 44: W242–W245. DOI:10.1093/nar/gkw290

46. Letzel A, Pidot S, Hertweck C. (2014). Genome mining for ribosomally synthesized and post-translationally modified peptides (RiPPs) in anaerobic bacteria. BMC Genomics 15: 983. https://doi.org/10.1186/1471-2164-15-983

47. Marchler-Bauer A, Lu S, Anderson J, Chitsaz F, Derbyshire M, DeWeese-Scott C, Fong JH, Geer LY, Geer RC, Gonzales NR, Gwadz M, Hurwitz DI, Jackson JD, Ke Z, Lanczycki CJ, Lu F, Marchler GH, Mullokandov M, Omelchenko MV, Robertson CL, Song JS, Thanki N, Yamashita RA, Zhang D, Zhang N, Zheng C, Bryant SH. (2010) CDD: a Conserved Domain Database for the functional annotation of proteins. Nucleic Acids Res 39: D225–D229. DOI:10.1093/nar/gkq1189

48. Maricic N, Anderson E, Opipari A, Yu E, Dawid S. (2016) Characterization of a multipeptide lantibiotic locus in *Streptococcus pneumoniae*. mBio 7: e01656–15. DOI:10.1128/mBio.01656-15

49. Martinez B, Rodriguez A, Suárez J, Fernández M. (1999) Synthesis of lactococcin 972, a bacteriocin produced by *Lactococcus lactis* IPLA 972, depends on the expression of a plasmid – encoded bicistronic operon. Microbiology 145: 3155–3161. DOI:10.1099/00221287-145-11-3155

50. Maynard Smith J, Dowson CG, Spratt BG. (1991) Localized sex in bacteria. Nature 349: 29–31. DOI:10.1038/349029a0

51. O’Brien K, Wolfson L, Watt J, Henkle E, Deloria-Knoll M, McCall N, Lee E, Mulholland K, Levine OS, Cherian T; Hib and Pneumococcal Global Burden of Disease Study Team.. (2009) Burden of disease caused by Streptococcus pneumoniae in children younger than 5 years: global estimates. Lancet 374: 893–902. DOI:10.1016/S0140-6736(09)61204-6.

52. Page A, Cummins C, Hunt M, Wong V, Reuter S, Holden MT, Fookes M, Falush D, Keane JA, Parkhill J. (2015) Roary: rapid large-scale prokaryote pan genome analysis. Bioinformatics 31: 3691–3693. DOI:10.1093/bioinformatics/btv421

53. Perez-Pascual D, Monnet V, Gardan R. (2016) Bacterial cell–cell communication in the host via RRNPP peptide-binding regulators. Front Microbiol 7: 706. DOI:10.3389/fmicb.2016.00706

54. Price M, Dehal P, Arkin A. (2010) FastTree 2 – approximately maximum-likelihood trees for large alignments. PLOS ONE 5: e9490. DOI:10.1371/journal.pone.0009490

55. Reichmann P, Hakenbeck R. (2000) Allelic variation in a peptide-inducible two-component system of *Streptococcus pneumoniae*. FEMS Microbiol Lett 190: 231–236. https://doi.org/10.1111/j.1574-6968.2000.tb09291.x

56. Seemann T. (2014) Prokka: rapid prokaryotic genome annotation. Bioinformatics 30: 2068–2069. DOI:10.1093/bioinformatics/btu153

57. Shak J, Vidal J, Klugman K. (2013) Influence of bacterial interactions on pneumococcal colonization of the nasopharynx. Trends Microbiol 21: 129–135. https://doi.org/10.1016/j.tim.2012.11.005

58. Son M, Shchepetov M, Adrian P, Madhi S, de Gouveia L, von Gottberg A, Klugman KP, Weiser JN, Dawid S. (2011) Conserved mutations in the pneumococcal bacteriocin transporter gene, *blpA*, result in a complex population consisting of producers and cheaters. mBio 2: e00179–11-e00179-11. DOI:10.1128/mBio.00179-11

59. Szklarczyk D, Franceschini A, Wyder S, Forslund K, Heller D, Huerta-Cepas J, Simonovic M, Roth A, Santos A, Tsafou KP, Kuhn M, Bork P, Jensen LJ, von Mering C. (2015) STRING v10: protein-protein interaction networks, integrated over the tree of life. Nucleic Acids Res 43:D447–52. DOI:10.1093/nar/gku1003

60. Tadesse B, Ashley E, Ongarello S, Havumaki J, Wijegoonewardena M, Gonzalez IJ, Dittrich S. (2017) Antimicrobial resistance in Africa: a systematic review. BMC Infect Dis 17: 616. https://doi.org/10.1186/s12879-017-2713-1

61. van Heel AJ, de Jong A, Montalbán-López M, Kok J, Kuipers OP. (2013) BAGEL3: Automated identification of genes encoding bacteriocins and (non-)bactericidal posttranslationally modified peptides. Nucleic Acids Res 41:W448–53. DOI:10.1093/nar/gkt391

62. van Tonder A, Bray J, Roalfe L, White R, Zancolli M, Quirk SJ, Haraldsson G, Jolley KA, Maiden MC, Bentley SD, Haraldsson Á, Erlendsdóttir H, Kristinsson KG, Goldblatt D, Brueggemann AB. (2015) Genomics reveals the worldwide distribution of multidrug-resistant serotype 6E pneumococci. J Clin Microbiol 53: 2271–2285. DOI:10.1128/JCM.00744-15

63. van Tonder A, Mistry S, Bray J, Hill D, Cody A, Farmer CL, Klugman KP, von Gottberg A, Bentley SD, Parkhill J, Jolley KA, Maiden MC, Brueggemann AB. (2014) Defining the estimated core genome of bacterial populations using a Bayesian decision model. PLOS Comput Biol 10: e1003788. DOI:10.1371/journal.pcbi.1003788

64. Walker G, Heng N, Carne A, Tagg J, Wescombe P. (2016) Salivaricin E and abundant dextranase activity may contribute to the anti-cariogenic potential of the probiotic candidate *Streptococcus salivarius* JH. Microbiology 162: 476–486. DOI:10.1099/mic.0.000237

65. Weber T, Blin K, Duddela S, Krug D, Kim H, Bruccoleri R, Lee SY, Fischbach MA, Müller R, Wohlleben W, Breitling R, Takano E, Medema MH. (2015) antiSMASH 3.0—a comprehensive resource for the genome mining of biosynthetic gene clusters. Nucleic Acids Res 43: W237–W243. DOI:10.1093/nar/gkv437

66. Health Organization World. (2017) Global priority list of antibiotic-resistant bacteria to guide research, discovery, and development of new antibiotics. http://www.who.int/medicines/publications/global-priority-list-antibiotic-resistant-bacteria

67. Wyres K, Lambertsen L, Croucher N, McGee L, von Gottberg A, Liñares J, Jacobs MR, Kristinsson KG, Beall BW, Klugman KP, Parkhill J, Hakenbeck R, Bentley SD, Brueggemann AB. (2012) Pneumococcal capsular switching: a historical perspective. J Infect Dis 207: 439–449. https://doi.org/10.1093/infdis/jis703

68. Zerbino D, Birney E. (2008) Velvet: Algorithms for de novo short read assembly using de Bruijn graphs. Genome Res 18: 821–829. DOI:10.1101/gr.074492.107

